# SARS-CoV-2 Genomic Surveillance in Costa Rica: Evidence of a Divergent Population and an Increased Detection of a Spike T1117I Mutation

**DOI:** 10.1101/2020.12.21.423850

**Authors:** Jose Arturo Molina-Mora, Estela Cordero-Laurent, Adriana Godínez, Melany Calderón-Osorno, Hebleen Brenes, Claudio Soto-Garita, Cristian Pérez-Corrales, COINGESA-CR Consorcio Interinstitucional de Estudios Genómicos del SARS-CoV-2 Costa Rica, Jan Felix Drexler, Andres Moreira-Soto, Eugenia Corrales-Aguilar, Francisco Duarte-Martínez

## Abstract

Genome sequencing is a key strategy in the surveillance of SARS-CoV-2, the virus responsible for the COVID-19 pandemic. Latin America is the hardest hit region of the world, accumulating almost 20% of COVID-19 cases worldwide. Costa Rica was first exemplary for the region in its pandemic control, declaring a swift state of emergency on March 16^th^ that led to a low quantity of cases, until measures were lifted in early May. From the first detected case in March 6^th^ to December 31^st^ almost 170 000 cases have been reported in Costa Rica, 99.5% of them from May onwards. We analyzed the genomic variability during the SARS-CoV-2 pandemic in Costa Rica using 185 sequences, 52 from the first months of the pandemic, and 133 from the current wave.

Three GISAID clades (G, GH, and GR) and three PANGOLIN lineages (B.1, B.1.1, and B.1.291) are predominant, with phylogenetic relationships that are in line with the results of other Latin American countries, suggesting introduction and multiple re-introductions from other regions of the world. The whole-genome variant calling analysis identified a total of 283 distinct nucleotide variants. These correspond mostly to non-synonymous mutations (51.6%, 146) but 45.6% (129) corresponded to synonymous mutations. The 283 variants showed an expected power-law distribution: 190 single nucleotide mutations were identified in single sequences, only 16 single nucleotide mutations were found in >5% sequences, and only two mutations in >50% genomes. These mutations were distributed through the whole genome. However, 63.6% were present in ORF1ab, 11.7% in Spike gene and 10.6% in the Nucleocapsid gene. Additionally, the prevalence of worldwide-found variant D614G in the Spike (98.9% in Costa Rica), ORF8 L84S (1.1%) is similar to what is found elsewhere. Interestingly, the frequency of mutation T1117I in the Spike has increased during the current pandemic wave beginning in May 2020 in Costa Rica, reaching 29.2% detection in the full genome analyses in November 2020. This variant has been observed in less than 1% of the GISAID reported sequences worldwide in all the 2020. Structural modeling of the Spike protein with the T1117I mutation suggest a potential effect on the viral oligomerization needed for cell infection, but no differences with other genomes on transmissibility, severity nor vaccine effectiveness are predicted. Nevertheless, in-vitro experiments are required to support these *in-silico* findings. In conclusion, genome analyses of the SARS-CoV-2 sequences over the course of COVID-19 pandemic in Costa Rica suggest introduction of lineages from other countries as travel bans and measures were lifted, similar to results found in other studies, as well as an increase in the Spike-T1117I variant that needs to be monitored and studied in further analyses as part of the surveillance program during the pandemic.

## INTRODUCTION

COVID-19 (COronaVIrus Disease 2019) is an infectious disease caused by the SARS-CoV-2 virus. It was first described in late December 2019, in an outbreak of atypical pneumonia in the city of Wuhan, Hubei province, China (Lu, Stratton, & Tang, 2020). Until the end of 2020, this virus has resulted in more than 81 million confirmed cases and 1.8 million deaths worldwide (https://coronavirus.jhu.edu/map.html). Latin America is the hardest hit region of the world, accumulating almost 20% of COVID-19 cases worldwide (https://ourworldindata.org/coronavirus). The first confirmed case of COVID-19 in Costa Rica was reported in San José (capital city) on March 6th, 2020. Costa Rica was first exemplary for the region in its pandemic control, declaring a swift state of emergency on March 16th that led to a low quantity of cases, until measures were lifted in early May. 99.5% of the cases have been found from May onwards, thus adding up to form a second bigger wave. Until December 31^st^, 2020 (Figure 1-A-B), Costa Rica has confirmed 169 321 cases and 2 185 deaths (https://www.ministeriodesalud.go.cr/).

**Figure 1.**
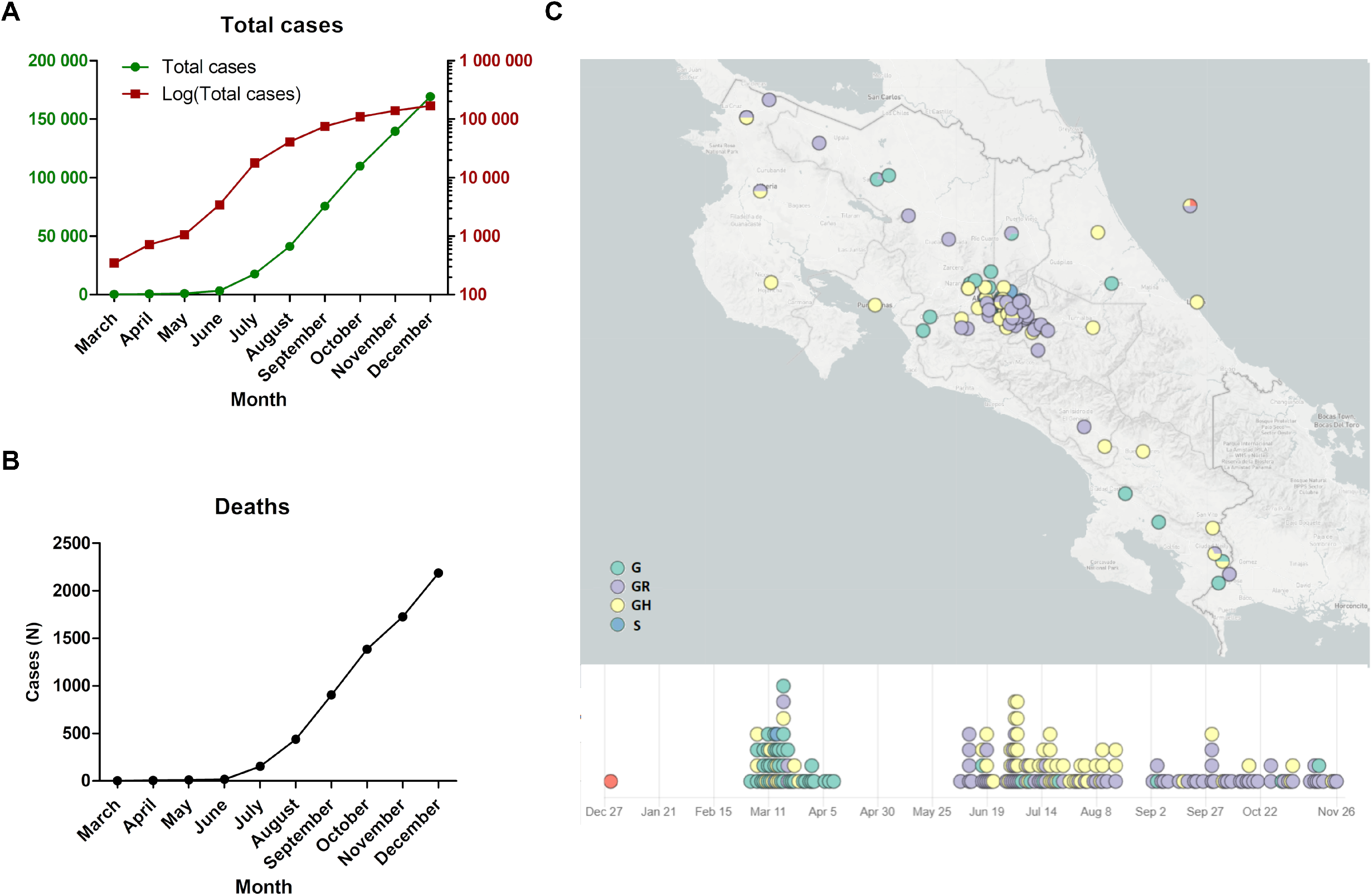
Dynamic, geographic and temporal distribution of SARS-CoV-2 genomes from Costa Rican cases. (A) An exponential increment of COVID-19 cases has been reported in Costa Rica since March 2020, with a similar profile for reported deaths (B). Samples for sequencing were obtained from the whole country, mainly from the Central valley, which harbors the most populated region of the country (C). Image in (C) was obtained from Microreact tool (https://microreact.org/project/r7tcnUYgWMRJ5Fdssvv7VZ).

The first complete SARS-CoV-2 genome globally was published on January 10, 2020, revealing that the genome is composed of a single-stranded RNA with 29 903 nucleotides (Wu et al., 2020). Genetic diversity of SARS-CoV-2 genomes has demonstrated a large number of independent introductions of the virus into many countries of the world but China (van Dorp et al., 2020). Although strong relationships between specific variants or clades in SARS-CoV-2 and the clinical outcome have not been identified (Grubaugh, Hanage, & Rasmussen, 2020; Hodcroft et al., 2020; van Dorp et al., 2020), different relevant parameters could be impacted by mutations in the virus genome. These include transmissibility, severity, clinical management, mortality, vaccines as well as the performance of molecular tests for diagnosis (Hodcroft et al., 2020; Koyama, Platt, & Parida, 2020; Teng, Sobitan, Rhoades, Liu, & Tang, 2020). Until now, host’s genetic factors and, more importantly, comorbidities seem to have the predominant impact on the fate of COVID-19 patients (LoPresti, Beck, Duggal, Cummings, & Solomon, 2020; Sironi et al., 2020; Toyoshima, Nemoto, Matsumoto, Nakamura, & Kiyotani, 2020). A few studies have suggested possible associations between genetic markers of the virus and clinical fate, which are still under investigation (Hodcroft et al., 2020; Korber et al., 2020; Toyoshima et al., 2020).

Due to new mutations and appearance over time of new genome clusters, whole-genome sequencing is a key step in the surveillance of this pathogen (Castillo et al., 2020). In this regard, diverse classification schemes using specific genetic markers have been proposed to monitor the evolution of the virus in the human population. The GISAID (Global Initiative on Sharing All Influenza Data, https://www.gisaid.org/) classification scheme by clades (GISAID, 2020) and the PANGOLIN (Phylogenetic Assignment of Named Global Outbreak LINeages) classification scheme by lineages (Rambaut et al., 2020) have been widely used for monitoring the current pandemic.

At the end of April 2020, genome sequencing of SARS-CoV-2 virus from Costa Rican cases of COVID-19 was started as part of the national epidemiological surveillance (INCIENSA & Ministerio de Salud, 2020). In this work we present the genome analysis of 185 SARS-CoV-2 sequences which have been obtained from Costa Rican cases of COVID-19 in the period March-November 2020, including 52 genomes from the first months of the pandemic, and 133 from the current second wave (from May onwards). The study aimed to investigate the diversity of the viral population of SARS-CoV-2 genomes circulating in Costa Rica. To this end, we included the genome sequencing and assembly, the analysis of variants or mutations, the construction of a phylogenetic tree, as well as the analysis of the possible association of genomes by clades/lineages with epidemiological data. Also, we report an increase in local detection of a variant with a Spike mutation T1117I not broadly reported elsewhere.

## METHODS

### Clinical Isolates

Material from nasopharyngeal swabs was obtained from Costa Rican cases of COVID-19, in the period between March and November 2020. All the patients were diagnosed in INCIENSA (Instituto Costarricense de Investigación y Enseñanza en Nutrición y Salud) or different public and private clinical laboratories by real-time reverse transcription polymerase chain reaction (RT-PCR), using specific probe and primers according to the guidelines suggested by the Pan American Health Organization and the World Health Organization (PAHO, 2020) and the Ministry of Health of Costa Rica. Only samples with a CT<25 were considered for the genome sequencing. Samples (167) were sent to the sequencing service of INCIENSA for further analyses. Other 18 assembled sequences of Costa Rica were included (see below, ID: CRC-007-024), completing 185 genomes. Gender distribution of the samples was 112 male and 71 female patients (two cases without gender information), and the age ranged from 4 to 92 years old. Ten deaths related to COVID-19 were included in these analyses. More details in Table 1.

**Table 1.** Demographic information and basic results of the genome analysis of SARS-CoV-2 from Costa Rican cases.

| Parameter | Counts |  |
| --- | --- | --- |
| Demographic data | ***Patients*** | ***Location (provinces)*** |
|  | Total cases: 185 | San José: 82 |
|  |  | Alajuela: 36 |
|  | ***Sex*** | Heredia: 16 |
|  | Male: 112 | Cartago: 20 |
|  | Female: 71 | Puntarenas: 13 |
|  | Unknown: 2 | Guanacaste: 9 |
|  |  | Limón: 6 |
|  | Mortality (known cases): 10 | Other: 3 |
| Genome sequencing | ***Genome sequencing*** | ***Genome coverage depth*** |
|  | Total genomes: 185 | 900-7000X |
| Total variants for all genomes | 2120, including repeated mutations among genomes |  |
| Total distinct variants | 283 variants, regardless frequency among genomes |  |
| Variability | Reference genome size: 29 903 bp |  |
|  | Variants (min-max) | 3-22 per genome |
|  | Identity: | 99.92 - 99.98% |
| Classification of variants | ***By gene*** | ***By effect*** |
|  | ORF1ab: 180 (63.6%) | Missense: 146 (51.6%) |
|  | ORF3a: 11 (3.9%) | Synonymous: 129 (45.6%) |
|  | ORF6: 1 (0.4%) | Splice region: 1 (0.3%) |
|  | ORF7a: 1 (0.4%) | STOP gained: 3 (1.1%) |
|  | ORF8: 6 (2.1%) | Upstream: 4 (1.4%) |
|  | S-Spike: 33 (11.7%) |  |
|  | E: 5 (1.8%) | ***By frequency*** |
|  | M: 9 (3.2%) | In only 1 genome: 190 (67.1%) |
|  | N: 30 (10.6%) | At least 5% (9) genomes: 16 (5.7%) |
|  | ORF10: 7 (2.5%) | At least 25% (46) genomes: 12 (4.2%) |
|  |  | At least 50% (92) genomes: 2 (0.7%) |
| Genome sequence groups: clades and lineages | ***GISAID clades*** | ***PANGOLIN lineages*** |
|  | G: 46 (24.9%) | A.1: 2 (1.1%) |
|  | GH: 60 (32.4%) | B.1: 28 (15.1%) |
|  | GR: 77 (41.6%) | B.1.1: 64 (34.6%) |
|  | S: 2 (1.1%) | B.1.203: 10 (5.4%) |
|  | V: 0 | B.1.291: 41 (22.2%) |
|  | L: 0 | B.1.5: 21 (11.3%) |
|  |  | Others: 19 (10.3%) |

### Genome sequencing

SARS-CoV-2 RNA was extracted with QIAamp Viral Mini Kit (Qiagen) in a QIACube extractor or with Maxwell^®^ RSC Viral Total Nucleic Acid Purification Kit (Promega) in a Maxwell^®^ RSC instrument following the manufacturer’s protocol. cDNA for library preparation was then synthesized and purified using either the ARTIC SARS-CoV-2 amplification protocol as described in https://artic.network/ncov-2019 (for samples 071-138), the SARS-CoV-2 amplification protocol described in (Castillo et al., 2020) (samples 001-006 and 026-070) or the FIOCRUZ protocol (samples 0140-0186) described in (Resende et al., 2020).

The genomics laboratory of INCIENSA prepared the sequencing libraries using the Illumina DNA Prep Kit (Illumina, San Diego, CA, USA) according to the laboratory standard operating procedure for pulsenet nextera DNA flex library preparation (https://www.cdc.gov/pulsenet/pathogens/wgs.html) as recommended by the Centers for Disease Control and Prevention (Atlanta, GA, USA). Paired-end sequencing (250 bp) was performed for each library on a MiSeq instrument using 500 cycles v2 chemistry cartridges (Illumina, San Diego, CA, USA).

### Genome assembly

Analyses were performed using custom scripts.

#### Quality control

Raw sequencing reads were evaluated using FastQC v0.11.7 (Andrews, 2010) and MultiQC (Ewels, Magnusson, Lundin, & Käller, 2016) before and after trimming. Removal of adapters and trimming of low-quality sequences (Q<30) was done Trimmomatic v0.38 (Bolger, Lohse, & Usadel, 2014). FastQ-Screen (Wingett & Andrews, 2018) and BBDuk (http://jgi.doe.gov/data-and-tools/bb-tools/) were used to quantify and remove possible contaminant reads, respectively.

#### Assembly strategies

Reference-based assembly (consensus sequence) was done by mapping reads to a reference SARS-CoV-2 genome NC_045512.2 using BWA-MEM 0.7.5a-r405 (H. Li & Durbin, 2009) with default parameters. The resulting BAM file was used for the variant calling analyses. In addition, SARS-CoV-2 *de novo* genome assembly was performed with Megahit v1.1.3 (D. Li, Liu, Luo, Sadakane, & Lam, 2015). Due to the results of both *de novo* and reference-based assemblies were consistent, the consensus sequence was used for subsequent analyses.

#### Assembly assessment

Assessment of genome assembly was done using the 3C criterion (contiguity, completeness, and correctness) as defined in (J.-A. Molina-Mora, Campos-Sánchez, Rodríguez, Shi, & García, 2020; J. A. Molina-Mora & Garcia, 2020). QUAST (Gurevich, Saveliev, Vyahhi, & Tesler, 2013), Qualimap (Okonechnikov, Conesa, & García-Alcalde, 2016), and BRIG (Alikhan, Petty, Ben Zakour, & Beatson, 2011) results were used for the analysis.

#### Clade assignment

PANGOLIN Lineages and GISAID clades were assigned based on assembled genomes using Bionumerics software v7.6.3 (https://www.applied-maths.com/) or the Coronavirus typing tool v1.13 (https://www.genomedetective.com/app/typingtool/cov/) after their upload into the GISAID database (Global Initiative on Sharing All Influenza Data, www.gisaid.org). Microreact (Argimón et al., 2016) was used to visualize each case genome according to geographic location and the cluster it belongs to. In addition, 18 assembled SARS-CoV-2 genomes from Costa Rica (CRC-007 to CRC-024, University of Costa Rica) were retrieved from the GISAID database and included for the comparative analyses.

### Comparative genomic analyses

#### Genome variability

Cd-hit (W. Li & Godzik, 2006) was used to cluster and compare all genome sequences by similarity. We discarded genome sequences belonging to other clusters than the one containing the reference. All the viral sequences passing the quality control step and which resulted clustered with the reference were uploaded into the GISAID database.

#### Variant calling analysis

BAM file from reads mapping was used to remove duplicates using Picard Markduplicates (http://broadinstitute.github.io/picard). Freebayes v1.3.1 (Garrison & Marth, 2012) was used as a variant caller with the parameters:-p 1-q 20-m 60--min-coverage 10 –V. VCF_filter v3.2 (https://github.com/moskalenko/vcf_filter) was used to remove low-confidence variants. Variant annotation was achieved using SNPeff (Cingolani et al., 2012) with the annotation file of the reference SARS-CoV-2 genome NC_045512.2. To represent relevant variants, structural modeling (3D) of Spike protein was done using PDB (https://www.rcsb.org/).

#### Phylogenetic analysis

MAFFT v7.471 (Katoh, Misawa, Kuma, & Miyata, 2002) was used to align all genome sequences. Construction of the phylogenetic tree was done using IQ-TREE v1.6.12 (Minh et al., 2020), including ModelFinder (Kalyaanamoorthy, Minh, Wong, Von Haeseler, & Jermiin, 2017) to select the best nucleotide substitution model (using the Bayesian Information Criterion BIC, the best model was TN+F+I). The tree was visualized using iTOL tool v4 (Letunic & Bork, 2019), including information of clades and lineages (GISAID and PANGOLIN classification), key variants, epidemiological week, and geographic location (Costa Rica is administratively composed of seven provinces which are divided into 82 cantons).

## RESULTS

To study the SARS-CoV-2 genomes from Costa Rican cases of COVID-19, sequencing, genome assembly, and genomic comparison were performed. Samples were collected from different geographical regions from Costa Rica from March to November 2020 (Figure 1 and Table 1), with a predominance of cases from the Central Valley region (the most highly populated region). No particular features were recognized by geographic origin at the genomic level (see below).

Variant calling analysis identified a range of 3 to 22 variants per genome. In total, we identified 283 distinct genome variants with a frequency of 2120 among all genomes. The total of distinct genome variants includes 146 (51.6%) missense (non-synonymous) mutations and 129 (45.6%) synonymous mutations. More details in Table 1.

The distribution of all the 283 distinct variants among the genomes is shown in Figure 2-A. A power-law pattern was recognized, in which scarce mutations are present in most of the sequences, and so many variants are present in very few sequences (Figure 2-B). This means that only a few variants are present in multiple genomes, and the subsequent analysis can be focused on those variants. For example, only 16 variants were present in at least 5% (9) of the sequences, and were distributed along the genome (Figure 2-C and Table 2). Also, only two variants were found in >50% sequences, and most variants (190 sequences, 67.1%) were found in a single genome. The most frequent substitutions were D614G in the Spike and P4715L in the ORF1ab-RdRp both in 98.9% (183) of the genomes. Details and other cases are presented in Table 2.

**Table 2.** Variants of SARS-CoV-2 genomes observed in more than 5% (9) genomes from Costa Rican cases of COVID-19.

| Position genome | Ref | Alt | Gene | Position - cDNA | Position - protein | Type | Frequency (N) | Frequency (%) |
| --- | --- | --- | --- | --- | --- | --- | --- | --- |
| 14408 | C | T | ORF1ab - RdRp | c.14144C>T | p.Pro4715Leu (P4715L) | missense | 183 | 98.9 |
| 23403 | A | G | Spike - Surface glycoprotein | c.1841A>G | p.Asp614Gly (D614G) | missense | 183 | 98.9 |
| 28881 | G | A | N - Nucleocapsid phosphoprotein | c.608G>A | p.Arg203Lys (R203K) | missense | 77 | 41.6 |
| 28882 | G | A | N - Nucleocapsid phosphoprotein | c.609G>A | p.Arg203Arg (R203R) | synonymous | 77 | 41.6 |
| 28883 | G | C | N - Nucleocapsid phosphoprotein | c.610G>C | p.Gly204Arg (G204R) | missense | 77 | 41.6 |
| 1059 | C | T | ORF1ab - NSP2 | c.794C>T | p.Thr265Ile (T265I) | missense | 70 | 37.8 |
| 10360 | G | A | ORF1ab - 3C-like proteinase | c.10095G>A | p.Lys3365Lys (K3365K) | synonymous | 66 | 35.7 |
| **24912** | **C** | **T** | **Spike - Surface glycoprotein** | **c.3350C>T** | **p.Thr1117Ile (T1117I)** | **missense** | **54** | **29.2** |
| 28706 | C | T | N - Nucleocapsid phosphoprotein | c.433C>T | p.His145Tyr (H145Y) | missense | 54 | 29.2 |
| 29144 | C | T | N - Nucleocapsid phosphoprotein | c.871C>T | p.Leu291Leu (L291L) | synonymous | 54 | 29.2 |
| 8572 | G | T | ORF1ab - NSP4 | c.8307G>T | p.Trp2769Cys (W2769C) | missense | 52 | 28.1 |
| 10340 | C | T | ORF1ab - 3C-like proteinase | c.10075C>T | p.Pro3359Ser (P3359S) | missense | 52 | 28.1 |
| 2716 | C | T | ORF1ab - NSP2 | c.2451C>T | p.Gly817Gly (G817G) | synonymous | 44 | 23.8 |
| 17010 | C | T | ORF1ab - helicase | c.16746C>T | p.Ile5582Ile (I5582I) | synonymous | 44 | 23.8 |
| 7482 | C | T | ORF1ab - NSP3 | c.7217C>T | p.Ser2406Leu (S2406L) | missense | 43 | 23.2 |
| 26143 | G | A | ORF3a | c.751G>A | p.Gly251Ser (G251S) | missense | 12 | 6.5 |

**Figure 2.**
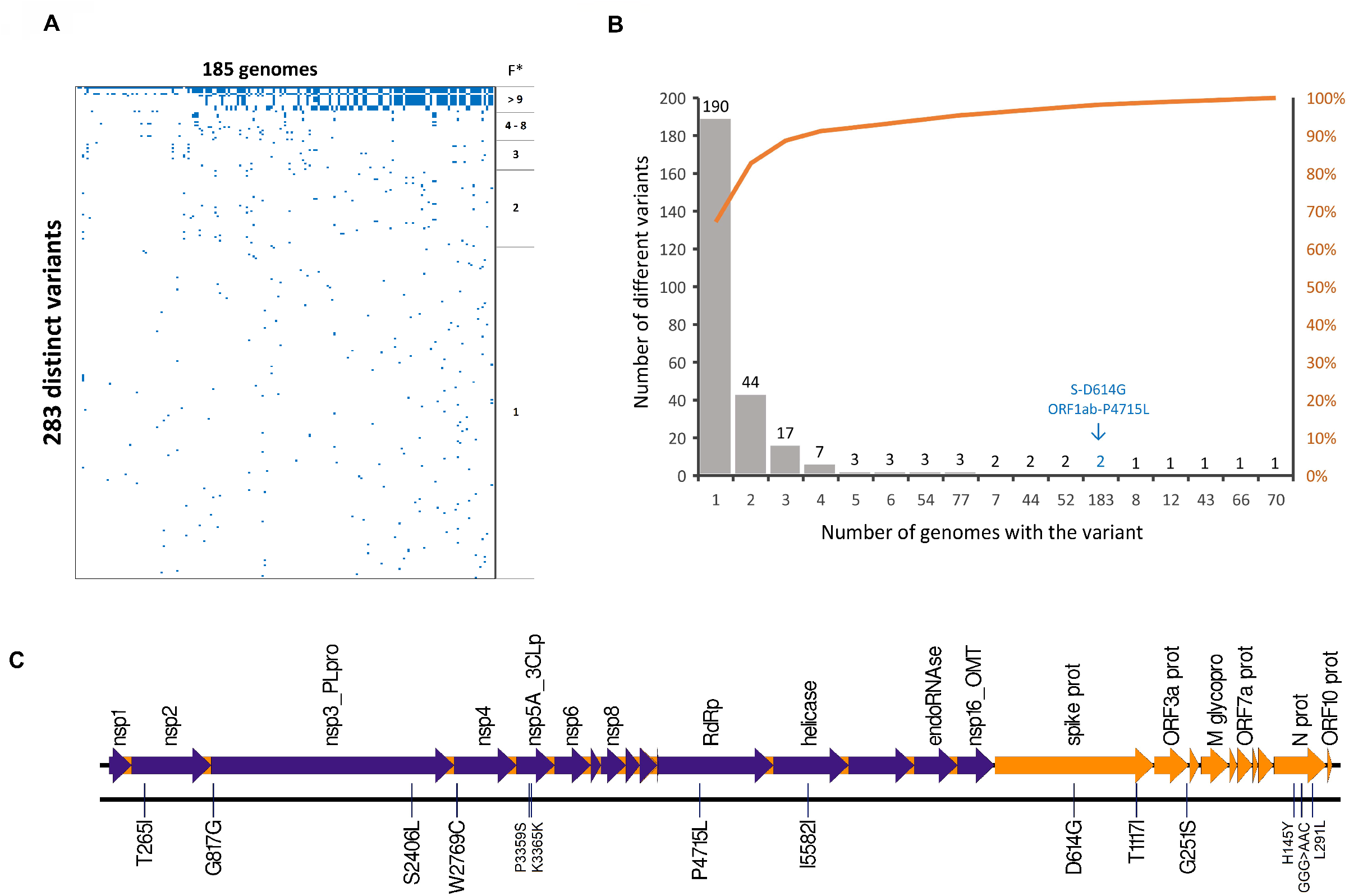
Variant calling analysis of SARS-CoV-2 genomes from Costa Rican cases of COVID-19. (A) Presence/absence of 283 different variants among 185 genomes. A few variants are widely distributed among genomes (*F= Frequency), and many variants are uniquely present in a single genome. (B) Distribution and accumulative percentage of variant frequency among genomes. Most variants are low frequency mutations and only 16 variants are present in at least 5% (9) genomes. The most frequent variants (Spike D614G and ORF1a P4715L) are present in 183 (98.9%) genomes (arrow). The 16 variants are distributed along the SARS-CoV-2 genome, as shown in (C).

Interestingly, the second most detected variant in the Spike gene in our analyses was the T1117I, present in 29.2% of the Costa Rican genomes. As shown in Figure 3, this mutation is very scarcely reported in the world (only present in 213 sequences, i.e., 0.08% according to GISAID, December 31^st^ 2020). In Costa Rica, the frequency during March, April, and May was 0%, in June reached 6.3% (5 out of 79 total sequences), July with 11.1 % (13 out of 117 genomes) and August with 14.5% with 20 sequences in 138 genomes. In September and October the frequency was 22.7 26.6%, respectively. To November, it reached 29.2% with 54 out of 185 genomes (Figure 3 and Supplementary Material). To better visualize the location and possible effects of this variant, a structural modeling of the Spike protein was done, as shown in Figure 4. The T1117I and D614G are located in different domains.

**Figure 3.**
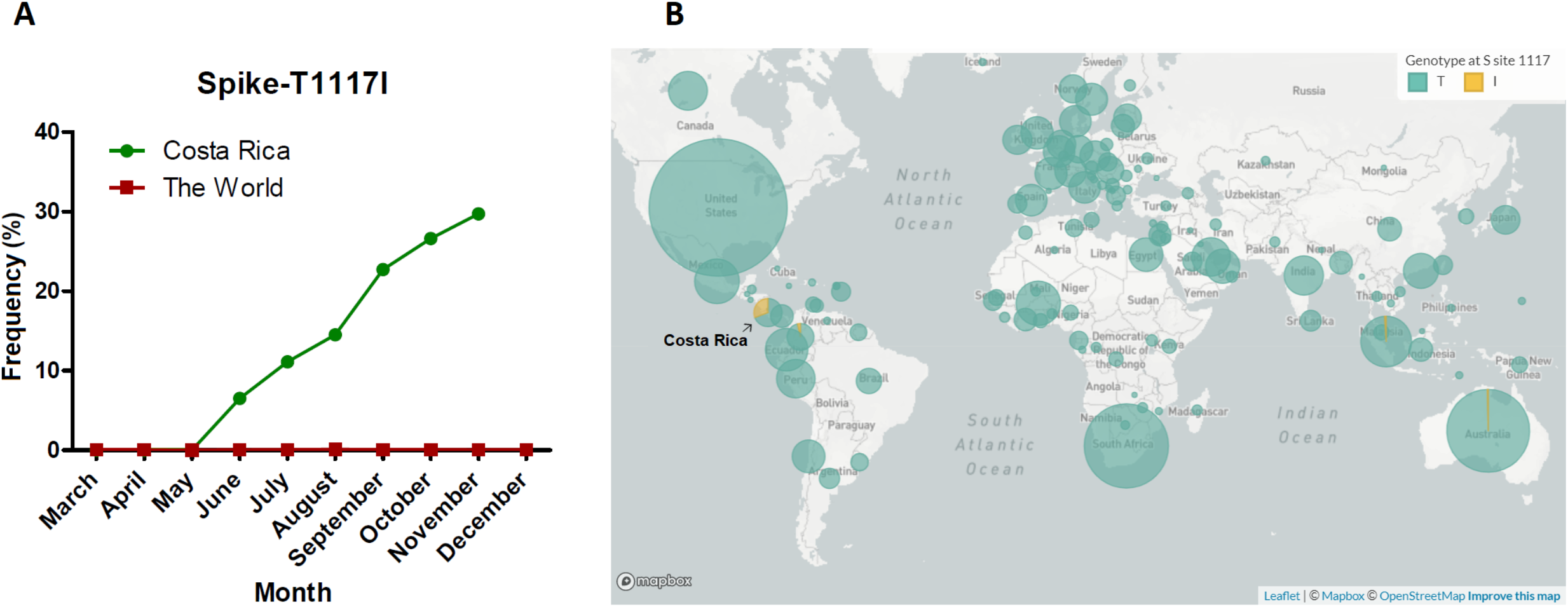
Frequency of T1117I in the spike along time in Costa Rica and the world. A notorious increment of the mutation has been reported in Costa Rica since May 2020, contrasting with the prevalence around the world which keeps relatively constant and low (A-B). Map in (B) was obtained from GISAID database.

**Figure 4.**
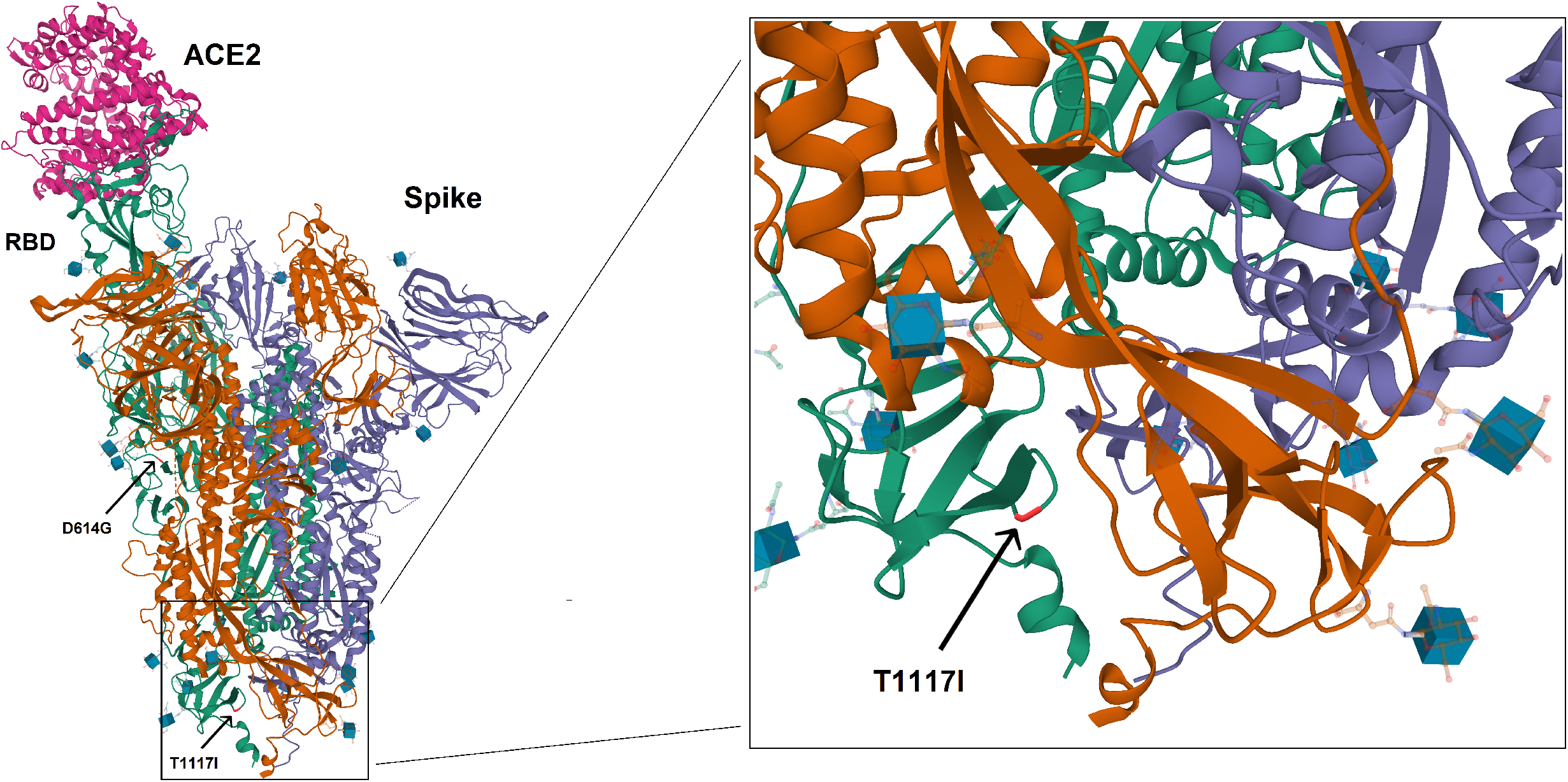
Structural modeling of spike protein of SARS-CoV-2. The variant D614G is present in 98.6% of the genomes in this study, which is also predominant worldwide (>90%, GISAID). D614G could affect the interaction with the host, as well as the immune response (vaccines), but real effects remains unclear. The variant T1117I is a variant very scarcely reported in the world (0.08%, GISAID), but the frequency in Costa Rica is 29.2%. The possible effect of this variant on the function of the spike is unknown.

On the other hand, the L84S variant in the ORF8 was only found in two genomes (the same without the D614G in the Spike). These two genomes were clustered separately as part of clade S in the phylogenetic tree (Figure 5, see details below).

**Figure 5.**
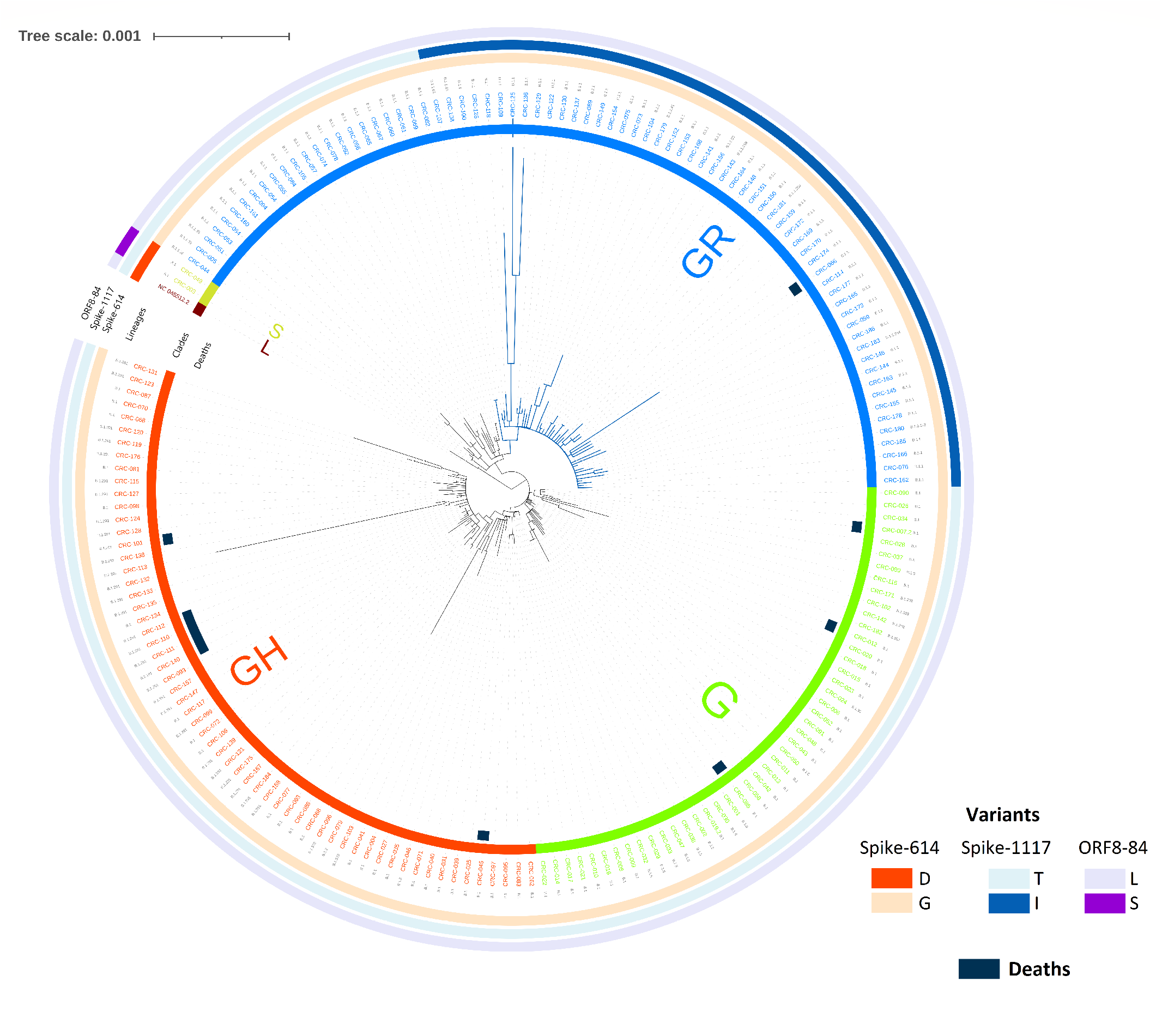
Phylogenetic tree of SARS-CoV-2 genomes circulating in Costa Rica. Three GISAID clades (G, GH and GR) and three Pangolin lineages (B.1, B.1.1 and B.1.291) are dominant in Costa Rican cases. Deaths are similarly distributed along clades. Variant T1117I in the spike is present in 54 genomes, which belong to a clearly separated monophyletic cluster (dark-blue). Other variants are presented with different colors.

In addition, no structural variants in the receptor-binding domain (RBD) of the Spike protein (N439K, T481I, V483A, E484E, N501Y nor G476S) were identified. Neither of the variants has been identified in the genomic regions that are used for the viral diagnosis by RT-PCR in the E and RdRP genes according to the protocol by Corman et al. 2020 (Corman et al., 2020).

Sequence classification showed that three major clades are predominant in Costa Rica according to both GISAID and PANGOLIN classifications (Table 1 and Figure 5). For GISAID clades, G, GH, and GR represent the main groups, with 98.9% of total cases, and two sequences were found as part of the S clade (1.1%). In the PANGOLIN classification, 20 different lineages were recognized, but 71.9% of the cases are represented by B.1, B.1.1, and B.1.291. No clear geographic distribution was found for the clades along the country (Figure 1 and Supplementary Material).

During the first five weeks of the outbreak in Costa Rica (from March to April) lineages B.1 and B.1.5 were mainly identify. The epidemiological information shown that those patients traveled to Europe (France and Spain), North America (USA and Mexico) and South America (Colombia). Later on lineage B.1.1 was documented on epidemiological week 24 and circulated actively in the north region of the country. Since then it was detected in samples of patients from six out of seven provinces from June to August 2020.

Based on the variants, the sequence comparison was done to build the phylogenetic tree, in which the best substitution model was TN+F+I (Figure 5). The clades and lineages are separated in the tree, including a clear cluster for all the 54 sequences carrying the Spike-T1117I variant (as part of the clade GR). Besides, the GR clade was enriched with cases from the 23-48 epidemiological weeks 2020 (June-November), but no other temporal patterns were identified. The only two sequences belonging to the clade S were clustered together and separately from the other Costa Rican cases, as expected. Regarding the ten deaths, four of the respective genomes clustered together (CRC-110, CRC-111, CRC-112 and CRC-140). Patients belonged to a nursing home outbreak. No epidemiological linkage was found between them and CRC-101. In general, fatal cases were distributed among the clades without a clear association with a specific taxonomic group.

## DISCUSSION

The analysis of genome sequences from pathogens has been recognized as a key tool in infectious disease epidemiology (van Dorp et al., 2020). Thus, to effectively combat COVID-19, epidemiological surveillance is a requirement to track circulating clades (Koyama et al., 2020).

In order to study the SARS-CoV-2 genome diversity circulating in Costa Rica, we compared 185 viral genomes. Concerning the variant calling analysis, the genome comparison against the reference genome (Wuhan-1, NC_045512.2) revealed between 3 and 22 mutations per genome, with a sequence homologies of 99.92-99.98%. Although SARS-CoV-2 is a RNA virus, the proof-reading activity of a nonstructural gene explains the relatively slow mutation rate of 1-2 mutations per month along the genome sequence (Deng et al., 2020). Interestingly, this observation is in line with our results, in which the genomes circulating in March-April 2020 reached up to 11 mutations, up to 18 in August, and up 22 in November.

In the analysis among all the genomes, 2120 mutations were identified, corresponding to 283 distinct genome variants. The last were represented mainly by missense (51.6%) and synonymous (45.6%) mutations. Similar distributions have been reported in other studies, in which the frequency of missense mutations in SARS-CoV-2 genomes is greater than synonymous mutations (Jaroszewski, Iyer, Alisoltani, Sedova, & Godzik, 2020; Koyama et al., 2020). By gene, most of the variants were identified in ORF1ab (63.6%), Spike (11.7%), and Nucleocapsid (10.6%). This predominance has also been reported in other studies (Vankadari, 2020).

Besides, the frequency of the distinct variants follows a power-law distribution. Only 16 out of the 283 variants were found in >5% of the genomes and most variants (67.1%) were found in a single genome. The most common variants were P4715L in the ORF1ab-RdRp and D614G in the Spike protein (both in the same 183 genomes, 98.9%). Similar findings have been reported in previous studies, as shown below.

### Spike protein - D614G

Since the Spike glycoprotein is involved in the interaction with the host’s receptor (angiotensin-converting enzyme 2, ACE2) (Zhou et al., 2020), variants in this gene are interesting because of the possible change in the dynamics of the transmission (Toyoshima et al., 2020) or vaccine effectiveness (Koyama et al., 2020). As observed also in Costa Rica (98.9%), D614G of the Spike protein is highly prevalent in the world, including America and Europe (Coppée, Lechien, Declèves, Tafforeau, & Saussez, 2020; GISAID, 2020; Mercatelli & Giorgi, 2020). According to GISAID, D614G already occurred worldwide in >90% of all samples.

Despite the relation between D614G and transmissibility, severity or mortality remains unclear (Grubaugh et al., 2020). Different studies have suggested some implications in transmission and immune response (Teng et al., 2020; Zhang et al., 2020), but analytical and epidemiological factors could explain this observation. In patients, D614G has been associated with higher viral load, but without significant changes of the disease severity (Korber et al., 2020; Plante et al., 2020) or vaccine effectiveness (Plante et al., 2020; Zhang et al., 2020).

### Spike protein - T1117I

T1117I in the Spike is the second variant found in frequency in the Spike protein (54 genomes, 29.2%). In our study, this mutation co-occurs with the H145Y and L291L variants in the Nucleocapsid. Interestingly, T1117I has been scarcely reported in other latitudes (GISAID, 2020; Long et al., 2020). According to GISAID (December 31^st^, 2020), T1117I occurred merely 213 times worldwide (0.08% of all samples with Spike sequence in the database) in only 19 countries, including Colombia and New Zealand. Our 54 sequences represent 25.3% out of all the available sequences with the T1117I variant worldwide. The group defines a clearly separated monophyletic cluster (see below, Figure 5). The first strain detected with this mutation worldwide was collected in March 2020 in Germany. In Costa Rica, T1117I was first reported in June 2020 in San José, and it increased in number during the June-December period (0%, 6.3%, 11.1%, 14.5%, 22.7%, 26.6% and 29.2% from May to November 2020, respectively), mainly in the Central Valley. Even though we cannot exclude a sampling bias from our data, the increasing detection of these variant through time should be further investigated using more sequences to reach a definite conclusion.

Using resolved structures of proteins from related strains, the Spike position equivalent to this mutation is involved in viral oligomerization interfaces needed for cell infection (Meher, Bhattacharjya, & Chakraborty, 2019; Walls et al., 2017; Wrapp et al., 2020). The radical replacement of threonine (T, a polar amino acid) by Isoleucine (I, non-polar) may have an impact on the Spike function. Due to the location of the T1117I variant in the Spike (Figure 3), it is not expected to induce direct changes in the interaction of the Spike (RBD) with the ACE2, nor the immunodominant region of the B-cell epitope (immune response) (Koyama et al., 2020). However, further studies are required to study the possible implications of this variant in the biology of SARS-CoV-2 and the spread in Costa Rica and the world.

### ORF1ab - P4715L (RdRp - P323L)

Other studies have shown that almost all sequences with the D614G (Spike) mutation also have the variant P4715L in the ORF1ab (RdRp, P323L). In Costa Rica, all the sequences with D614G also carry the P4715L, with 98.9% out of all the genomes included here. As part of the RdRp, P4715L might affect the replication speed of the virus (Koyama et al., 2020). The relevance of variants in the non-structural proteins is related to functions related to viral replication and transcription, including genes such as RdRp (Toyoshima et al., 2020). Also, because RdRp is the target of putative anti-viral drugs (such as remdesivir and favipiravir), mutations in these regions could eventually be responsible for the emergence of drug-resistant virus (Koyama et al., 2020). However, as found for other variants, the functional significance of P4715L and other variants in the RdRp is yet unknown (Toyoshima et al., 2020).

### ORF8-L84S

Only two genomes (1.1%) in this study have the variant L84S in the ORF8. L84S, one out of the two variants that define the clade S (GISAID classification), has been reported mainly for Asian countries, with a lower frequency in European and American countries (Coppée et al., 2020). This mutation was also implicated in the initial appreciation of the L and S genome types (Tang et al., 2020). However, like the other variants, this condition remains an open problem, in part due to the loss of the patterns as the available genome sequences have increased.

### Nucleocapsid, GGG>AAC

In the case of the variation GGG>AAC in the positions 28881-28883, the triplet resulted present in 77 (41.6%) genomes. This complex affects the Nucleocapsid protein with two amino acid changes and it has been reported in multiple latitudes (GISAID, 2020; Maitra et al., 2020; Osório & Correia-Neves, 2020). This complex variation was reported at the beginning of the binding site of the forward primer used in the RT-qPCR assays of the Chinese National Institute for Viral Disease Control and Prevention (Osório & Correia-Neves, 2020), however, this is not part of the primers used in other protocols, including those used in this study or any Costa Rican diagnostic laboratories.

### Other variants

Recently, a variant N501Y in the Spike has been reported in the United Kingdom, which is located in the RBD and the impact on the transmission or severity is not known (Kemp et al., 2020; Wise, 2020). From the Costa Rican sequenced cases, none harbors yet this mutation, neither the variant 501Y.V2 described in South Africa (Tegally et al., 2020). Also, a European cluster (called 20A.EU1) has been reported with a significant spread in summer 2020, mainly in Spain (Hodcroft et al., 2020). Genome sequences in this group have at least six variants, including the mutation A222V in the Spike protein and A220V in the nucleoprotein. None of the A222V in the Spike and the A220V in the nucleoprotein were found in our study. Nevertheless, local continued genomic surveillance is in place to monitor these variants.

Regarding the RT-qPCR test for the diagnosis of COVID-19, none of the 283 variants were part of the genomic regions of the binding site of the primers used in the Cormans et al. protocols (Corman et al., 2020) implemented in Costa Rica for detecting E and RdRp genes of SARS-CoV-2.

### Phylogeny and Integration

Altogether, the variant calling analysis reveals that the patterns found in Costa Rica, with 283 distinct mutations, are similar to those observed worldwide (such as D614G in the Spike and L84S in the ORF8), except for the T1117I in the Spike. In general, these mutations define the genome groups, in which three GISAID clades and three PANGOLIN lineages were found to predominate in Costa Rica. The 54 sequences with the T1117I variant defines a separated cluster with a monophyletic origin, i.e. they have a common ancestor.

The evolutionary analysis of the SARS-CoV-2 genomes from Costa Rican cases suggests the possibility of multiple introductions into the Costa Rican population, as it has been reported worldwide (Koyama et al., 2020; van Dorp et al., 2020). In addition, the sequences from the first months of the pandemic grouped in lineage B.1 and B.1.5 mainly, suggesting low undetected circulation and re-introductions of new lineages not detected in the country during early stages of the pandemic due to the extreme lockdown measures. Although travel records and some contact tracing data are available for many cases, the reconstruction of the possible routes for the spread of particular clades is difficult in this case due we only have sequenced 185 out of the 169 321 (to December 31^st^, 2020) Costa Rican cases of COVID-19 (0.1%).

Concerning deaths, genomes analyzed from ten dead patients were distributed among the clades without a clear association pattern for a specific group. This distribution is similar to that found around the world in which, until now, no mutation or clade has been clearly associated with a higher fatality rate (Grubaugh et al., 2020; Vankadari, 2020). In the phylogenetic three, a cluster with four sequences of dead patients can be highlighted. It was related to an outbreak occurred in a nursing home for the elderly.

Finally, limitations of our study include a low number of samples, the non-randomized sampling used for the selection of the samples (sampling bias), incomplete clinical and epidemiological data such as travel records and contact tracing for all sequenced samples, and lack of sequenced genome information from key neighboring countries with a strong immigration practice to Costa Rica. Nevertheless this work adds to the efforts of the global scientific community to produce and publicly share SARS-CoV-2 genome sequences. The analyses reported here contribute to the monitoring of the spread of SARS-CoV-2 as part of the surveillance programs during the pandemic, and suggest that surveillance should be continued and (financially-) supported to keep track of important changes in the SARS-CoV-2 genome.

## CONCLUSION

In summary, the phylogenetic and variant patterns are similar to those observed worldwide, such as the high prevalence of variant D614G in the Spike and low frequency of L84S in the ORF8, as well as the arrangement by clades. Interestingly, the variant T1117I in the Spike has increased its frequency along time from March to November 2020. This contrasts with the frequency of 0.08% in the world, in which 54 (25.3%) out of all the only 213 sequences in the world are from Costa Rica. Taken together, at the beginning of the pandemic few lineages with few mutations circulated in our country. Restrictions were lifted and borders were opened therefore diversity of lineages and mutations increased. The Spike variant T1117I must be closely surveyed to see if it will become the predominant circulating variant in our country. Though this variant putative biological and clinical relevance is still unknown.

In addition, none of the variants was found in genomic regions used for diagnosis with the protocols implemented in Costa Rica. Until now, changes in severity, deaths, or transmission have not been recognized for specific mutations or clades of SARS-CoV-2 in the world. Nevertheless, further genome sequencing analyses are required to study the impact of the genomic features on the biology of the virus, transmissibility and spread, clinical outcome, and treatment/vaccine effectiveness.

## Acknowledgments

We are grateful to clinicians, microbiologists, and other personnel of public (Caja Costarricense de Seguro Social CCSS) and private clinical laboratories for the samples of confirmed cases of COVID-19. We also thank Carlos Mora Garro for his participation as assistant, and all members of CIET-University of Costa Rica, INCIENSA and CCSS for their logistic and financial support in the activities associated with the project.

## Author contributions

J.M.M., F.D.M., H.B. and C.S.G. participated in the conception and design of the study. H.B, C.S.G, C.P.C. and COINGESA-CR members were involved in sample processing for diagnosis of COVID-19. F.D.M., E.C.L., A.G and M.C.O. sequenced the genomes. J.M.M. implemented and standardized the bioinformatics pipelines. J.M.M., F.D.M., A.M.S., E.C.A. and J.F.D. participated in the interpretation of results. J.M.M. drafted the manuscript. All authors reviewed and approved the final manuscript.

## Ethical approval and consent to participate

Not applicable.

## Consent for publication

Not applicable.

## Availability of data and material

Genome sequences were uploaded to the GISAID database. All the IDs and related information are available in the “Supplementary Material”.

## LEGENDS OF FIGURES AND TABLES

**Supplementary Material.xlsx**

